# ESCRT Machinery Dysfunction in Motor Neurone Disease: TSG101, CHMP2B, and VPS4a Differentially Regulate TDP-43 Pathology, Autophagy, and Exosome Biogenesis

**DOI:** 10.64898/2026.07.01.735805

**Authors:** Laila Mohamed, Mohamed Fouad Shalaby, Ritchie Williamson, Samantha Louise McLean, Sriharsha Kantamneni

## Abstract

Motor Neurone Disease (MND) is characterised by progressive degeneration of upper and lower motor neurons, accompanied by cytoplasmic mislocalisation and hyperphosphorylation of TDP-43 — hallmarks that implicate failure of endolysosomal proteostasis. The Endosomal Sorting Complexes Required for Transport (ESCRT) pathway governs multivesicular body (MVB) formation, lysosomal cargo delivery, and autophagosome closure, yet its expression profile in human MND tissue and mechanistic contribution to disease pathology have not been established. Here, we report subunit-specific dysregulation of ESCRT proteins in postmortem motor cortex and spinal cord from MND patients: CHMP2B (ESCRT-III) is significantly upregulated in both regions, whilst TSG101 (ESCRT-I) and VPS37A (ESCRT-I) are significantly downregulated in motor cortex, indicating a region-specific remodelling of the ESCRT network. In a tunicamycin-induced ER stress model using NSC-34 motor neuron-like cells and primary cortical neurons, TSG101 overexpression reduced total and phosphorylated TDP-43, suppressed mTOR signalling, and restored autophagic flux, whereas TSG101 knockdown exacerbated TDP-43 accumulation and cytoplasmic mislocalisation. CHMP2B modulation selectively regulated TDP-43 phosphorylation without altering total TDP-43 levels, consistent with a casein kinase 1-dependent mechanism operating independently of bulk autophagy. Both TSG101 and VPS4a were required to maintain neuronal CD9 tetraspanin localisation to early endosomes; their depletion redirected CD9 to late endosomal and lysosomal compartments under ER stress. Extracellular vesicle characterisation revealed a functional divergence: TSG101 is required for general exosome biogenesis, whereas VPS4a ATPase activity specifically mediates loading of pathological TDP-43 cargo into EVs. Dynamic light scattering confirmed that ER stress and ESCRT modulation produce distinct, condition-specific alterations in EV size and polydispersity. These findings establish ESCRT dysfunction as a multifaceted contributor to MND pathogenesis and identify TSG101, CHMP2B, and VPS4a as mechanistically distinct therapeutic targets warranting preclinical validation.

## Introduction

Motor Neuron Disease (MND) is an adult-onset, progressive, and ultimately fatal neurodegenerative disorder caused by the selective degeneration of upper motor neurons in the motor cortex and lower motor neurons in the brainstem and spinal cord [1]. Clinically, this manifests as progressive muscle weakness, atrophy, fasciculations, and spasticity, leading to paralysis and death, typically within 3–5 years of symptom onset [2]. The global incidence of MND is approximately 1–3 per 100,000 person-years, with a lifetime risk of 1 in 300–400 [3]. While the majority (90–95%) of cases are sporadic with no clear genetic aetiology, approximately 5–10% are familial, often linked to mutations in genes such as chromosome 9 open reading frame 72 (C9orf72), superoxide dismutase 1 (SOD1), transactive response DNA-binding protein (TARDBP), and fused in sarcoma (FUS) [4, 5]. Despite advances in understanding disease mechanisms, current treatments remain limited, with only two disease-modifying therapies approved—riluzole and edaravone—offering modest survival benefits without halting neurodegeneration [6]. There is therefore an urgent unmet clinical need to identify novel therapeutic targets applicable to both familial and sporadic forms of MND.

A unifying pathological hallmark in the majority of MND cases, regardless of genetic background, is the mislocalisation and aggregation of the nuclear RNA-binding protein TAR DNA-binding protein 43 (TDP-43) [7, 8]. In healthy neurons, TDP-43 is predominantly nuclear and functions in RNA processing, including splicing, transport, and stabilisation [9]. In MND, TDP-43 is cleared from the nucleus and forms hyperphosphorylated, ubiquitinated, and cleaved cytoplasmic aggregates that are thought to exert toxic gain-of-function effects while also causing loss of normal nuclear function [10, 11]. Clearance of these aggregates is predominantly mediated by the ubiquitin-proteasome system (UPS) and the autophagy-lysosome pathway; however, both are progressively overwhelmed in MND, perpetuating a cycle of proteostatic failure [12]. Disruption of autophagic flux and endolysosomal trafficking are now recognised as early and mechanistically central events in MND pathogenesis [13, 14], and the molecular machinery governing these pathways represents a tractable target for therapeutic intervention.

The Endosomal Sorting Complexes Required for Transport (ESCRT) pathway is the principal molecular effector of endolysosomal cargo sorting and membrane remodelling in eukaryotic cells. [15, 16]. The ESCRT system comprises several multiprotein complexes (ESCRT-0,-I,-II,-III) and associated factors such as ALG-2-interacting protein X (ALIX) and the AAA-ATPase VPS4, which function sequentially to recognise ubiquitinated membrane proteins, invaginate the endosomal membrane, and ultimately cleave intraluminal vesicles (ILVs) that contain cargo destined for lysosomal degradation [17, 18]. Key subunits of direct relevance to the present study include TSG101 (ESCRT-I), VPS37A and VPS37B (ESCRT-I), and CHMP2B (ESCRT-III). Beyond endosomal trafficking, ESCRT proteins also play critical roles in autophagosome closure, amphisome formation, and the biogenesis of extracellular vesicles (exosomes), derived vesicles that release both physiological and pathological protein cargo into the extracellular milieu [19, 20, 21]. The tetraspanins CD63 and CD9 are established markers of exosomal membranes and are sorted through ESCRT-dependent endosomal pathways; their trafficking is therefore sensitive to ESCRT integrity. Collectively, the ESCRT pathway sits at the convergence of protein degradation, autophagic flux, membrane protein homeostasis, and intercellular communication — processes each critically implicated in MND pathogenesis [19, 20, 21].

ESCRT dysfunction has been implicated in several neurodegenerative diseases, providing a mechanistic rationale for investigating its role in MND. For instance, mutations in CHMP2B, which encodes a core ESCRT-III component, cause familial frontotemporal dementia (FTD) and have been identified in MND patients [22, 23]. Similarly, altered expression of TSG101, an ESCRT-I subunit, has been linked to impaired clearance of amyloid precursor protein (APP) and tau in Alzheimer’s disease models [24, 25]. Furthermore, genome-wide association studies have identified variants in ESCRT-related genes as potential risk factors for neurodegenerative diseases [26]. However, the expression profiles of ESCRT components in human MND tissues and their mechanistic contributions to TDP-43 pathology, autophagy, and exosome biology are unknown.

In this study, we systematically investigated the expression of key ESCRT proteins such as TSG101, CHMP2B, VPS37A, VPS37B, ALIX, and signal transducing adaptor molecule 1 (STAM1) in postmortem motor cortex and spinal cord tissues from MND patients and age-matched controls. Using a tunicamycin-induced ER stress model in NSC-34 motor neuron-like cells and primary cortical neurons, and TDP-43 pathology, we further explored the functional consequences of modulating ESCRT components on TDP-43 aggregation, phosphorylation, and subcellular localisation, autophagy flux, mTOR signaling, and membrane protein stability. Additionally, we examined the roles of TSG101 and VPS4a in exosome biogenesis and in the selective loading of pathological TDP-43 cargo into EVs, and characterised EV physical properties by dynamic light scattering. Together, these studies establish ESCRT dysfunction as a mechanistically diverse contributor to MND pathogenesis and provide a framework for targeted therapeutic modulation of individual ESCRT components.

## Materials and Methods

### Human Tissue Samples

Human postmortem motor cortex and spinal cord tissues were obtained from the London Neurodegenerative Diseases Brain Bank at the Institute of Psychiatry, Psychology and Neuroscience, King’s College London (London, UK). Ethical approval was granted by the relevant institutional review boards, REC Ref no18/WA/0206. Tissues were collected from 12 MND patients and 12 age-matched neurologically healthy controls. For each region, 6 samples per group were used for western blot analysis, a sample size consistent with previous studies in the field (57, 58). Tissues were stored at −80 °C until processing.

### Tissue Lysate Preparation

Frozen tissue samples were homogenised in ice-cold lysis buffer (50 mM Tris-HCl pH 7.4, 150 mM NaCl, 1 mM EDTA, 1% Triton X-100, 0.1% SDS) supplied with protease inhibitor cocktail, and phosphatase inhibitors (Thermo Fisher Scientific, UK). Homogenates were incubated on ice for 30 min, sonicated briefly (two 5-s pulses at 30% amplitude), and centrifuged at 12,000 × g for 20 min at 4 °C. Supernatants were collected, and protein concentrations were determined using the BCA assay (Thermo Fisher Scientific, UK). All samples were normalised to equal protein concentrations prior to downstream analysis.

## Cell Culture

### NSC-34 cell line

The mouse NSC-34 motor neuron-like cells (Cedarlane, Canada) were cultured in Dulbecco’s Modified Eagle Medium (DMEM; Gibco, UK) supplemented with 10% fetal bovine serum (FBS; Gibco, UK) and 1% penicillin-streptomycin (PNS; Gibco, UK) at 37 °C in a humidified atmosphere containing 5% CO₂. Cells were maintained at sub-confluent densities and used up to passage 30.

For differentiation, cells were seeded at 5 × 10⁵ cells/well in 6-well plates and allowed to adhere for 24 h. Differentiation was induced by replacing the medium with DMEM/F-12 (Gibco) supplemented with 1% FBS, 1% PNS, and 1 µM all-trans retinoic acid (Sigma-Aldrich, UK) for 5–7 days, with medium changes every 2 days.

### Primary neuronal cells

All procedures were performed in accordance with the UK Animals (Scientific Procedures) Act 1986 and approved by the University of Bradford Animal Welfare and Ethical Review Body. Primary cortical and hippocampal neurons were obtained from embryonic day 18 (E18) Wistar rat embryos (Charles River, UK) and cultured as described previously.

Brains were dissected in ice-cold Hanks’ Balanced Salt Solution (HBSS) without calcium and magnesium (Gibco, UK). Meninges, cerebellum, and optic nerves were removed, and the cortex and hippocampus were isolated. Tissue was chopped into small pieces and washed three times with HBSS. Enzymatic dissociation was performed using 0.05% trypsin-EDTA (Gibco, Thermo Fisher Scientific, Paisley, UK) for 15 min at 37 °C, followed by addition of DNase I (20 µg/mL; Sigma-Aldrich, UK) for 5 min. Tissue was gently triturated in HBSS containing 10% FBS to inactivate trypsin. Cells were pelleted, resuspended in Neurobasal medium (Gibco, UK) supplemented with B-27 (Gibco, Thermo Fisher Scientific, Paisley, UK), GlutaMAX (Gibco, UK), and 1% PNS, and plated onto poly-L-lysine (0.1 mg/mL; Sigma-Aldrich, UK)-coated coverslips or culture dishes. Cells were maintained at 37 °C in 5% CO₂, with half-media changes every 3–4 days. Viral transduction was done 7–10 days in vitro (DIV) and experiments were performed at 13–17 DIV.

### ER Stress Induction

Tunicamycin (Sigma-Aldrich, UK) was dissolved in DMSO as a 1 mM stock solution and stored at −20 °C. Differentiated NSC-34 cells or primary neurons were treated with 100 nM tunicamycin for 48 h to induce endoplasmic reticulum stress. Control cells received an equivalent volume of vehicle (DMSO, final concentration ≤0.1%).

## Plasmid Constructs and Transfections

### NSC-34 Transfections

The following plasmids were used: pCMV6-TSG101 (Origene, RC204500), pCMV6-CHMP2B (Origene, RC210800), pCMV6-empty (P.CAT) as control, TSG101 shRNA (Origene, TG511575), CHMP2B shRNA (Origene, TG511574), and scrambled shRNA (Origene, TR30012). Transfections were performed using Lipofectamine 2000 (Invitrogen, UK) according to the manufacturer’s protocol. Briefly, cells at 70–80% confluence were transfected with 2 µg plasmid DNA and 5 µL Lipofectamine 2000 in Opti-MEM (Gibco). Medium was replaced after 6 h, and cells were harvested or treated 48 h post-transfection.

### Primary Neuron Lentiviral Transduction

Lentiviral particles were produced using a second-generation packaging system. HEK293T cells (ATCC, CRL-3216) were cultured in DMEM supplemented with 10% FBS and 1% penicillin-streptomycin. At 70–80% confluence, cells were co-transfected with the target lentiviral vector and the packaging plasmids pLP1, pLP2, and pVSV-G (Thermo Scientific, UK) using Lipofectamine 3000 (Invitrogen, UK) according to the manufacturer’s protocol. Briefly, a total of 15 µg DNA (target vector:packaging plasmids at a 1:1:1:1 ratio) was diluted in Opti-MEM (Gibco) with P3000 reagent, mixed with Lipofectamine 3000, and added dropwise to HEK293T cells. Culture supernatants containing lentiviral particles were collected 48 h and 72 h post-transfection, pooled, and filtered through 0.45 µm low-protein-binding filters (Millipore). Viral particles were concentrated by ultracentrifugation at 50,000 × g for 2 h at 4 °C using a Beckman Optima XE-90 ultracentrifuge with SW32 Ti rotor. Viral pellets were resuspended in sterile PBS, aliquoted, and stored at −80 °C until use. Viral titers were determined by limiting dilution assay in HEK293T cells, with functional titers typically ranging from 1 × 10⁸ to 5 × 10⁸ infectious units/mL.

The following lentiviral constructs were used in this study, all generated by Vector Builder: TSG101 shRNA (Vector Builder, VB010101-9298) targeting the sequence 5’-GGAGCTGTGTTACTGCTTCAT-3’; a non-targeting control shRNA (SCR; Vector Builder, VB010102-0001) containing a scrambled sequence with no homology to any known mammalian gene; TSG101-WT (Vector Builder, VB010103-1234) encoding the full-length human TSG101 coding sequence; VPS4a-WT (Vector Builder, VB010104-5678) encoding the full-length human VPS4a coding sequence; and DN-VPS4a (Vector Builder, VB010105-9101) encoding human VPS4a with a glutamine substitution at residue 228 (E228Q), rendering the ATPase activity-deficient and conferring dominant-negative function.

For transduction of primary neurons, cells were plated at DIV 7 at a density of 2 × 10⁵ cells per well in 24-well plates or 5 × 10⁵ cells per well in 6-well plates containing poly-L-lysine-coated coverslips or culture dishes. Cells were transduced at a multiplicity of infection (MOI) of 10 in Neurobasal medium containing 5 µg/mL polybrene (Sigma-Aldrich) to enhance transduction efficiency. The medium was replaced after 24 h with fresh Neurobasal medium supplemented with B-27 (Gibco) and GlutaMAX (Gibco). Cells were maintained for an additional 72 h to allow for transgene expression before experimental treatments or harvesting.

Transduction efficiency was routinely assessed by quantifying EGFP or mCherry fluorescence using fluorescence microscopy (Nikon Eclipse Ti2) with automated image acquisition and analysis. Typical infection rates exceeded 70% across independent experiments. For experiments requiring uniform transgene expression, parallel wells were processed to confirm knockdown or overexpression efficiency by western blotting prior to experimental treatments.

### Protein Extraction

Cells were washed twice with ice-cold Tris-buffered saline (TBS; 50 mM Tris-HCl pH 7.4, 150 mM NaCl) and lysed in RIPA buffer (50 mM Tris-HCl pH 7.4, 150 mM NaCl, 1 mM EDTA, 1% Triton X-100, 0.5% sodium deoxycholate, 0.1% SDS) supplemented with protease inhibitor cocktail (Roche, UK) and phosphatase inhibitors (1 mM Na₃VO₄, 50 mM NaF, 5 mM sodium pyrophosphate). Lysates were incubated on ice for 30 min, sonicated, and centrifuged at 12,000 × g for 15 min at 4 °C. Supernatants were collected, and protein concentrations were determined using the BCA assay (Thermo Fisher Scientific).

### Western Blotting

Protein samples (20–40 µg per lane) were separated by SDS-PAGE on 4–20% or 10% polyacrylamide gels (Bio-Rad, UK) and transferred onto PVDF membranes (0.45 µm; Bio-Rad) using a wet transfer system (Bio-Rad). Membranes were blocked in 5% non-fat milk or 5% bovine serum albumin (BSA) in Tris-buffered saline containing 0.1% Tween-20 (TBST) for 1 h at room temperature. Primary antibodies were diluted in blocking buffer and incubated overnight at 4 °C with gentle agitation. The following primary antibodies were used: rabbit anti-TSG101 (1:1000; Abcam, ab125011), rabbit anti-CHMP2B (1:1000; Abcam, ab228841), rabbit anti-VPS37A (1:500; Proteintech, 16094-1-AP), rabbit anti-VPS37B (1:500; Proteintech, 25334-1-AP), mouse anti-ALIX (1:1000; Santa Cruz, sc-53538), rabbit anti-STAM1 (1:1000; Cell Signaling Technology, 12475), mouse anti-TDP-43 (1:1000; Proteintech, 60019-2-Ig), rabbit anti-phospho-TDP-43 (Ser409/410; 1:1000; Cosmo Bio, CAC-TIP-PTD-M01), rabbit anti-mTOR (1:1000; Cell Signaling Technology, 2983), rabbit anti-phospho-mTOR (Ser2448; 1:1000; Cell Signaling Technology, 5536), rabbit anti-LC3B (1:1000; Cell Signaling Technology, 2775), mouse anti-p62 (1:1000; BD Biosciences, 610832), mouse anti-β-actin (1:5000; Sigma-Aldrich, A5441), and rabbit anti-α-tubulin (1:5000; Abcam, ab18251). After washing with TBST, membranes were incubated with HRP-conjugated secondary antibodies (1:5000; Bio-Rad) for 1 h at room temperature. Immunoreactive bands were visualised using enhanced chemiluminescence (ECL; Bio-Rad) and imaged on a ChemiDoc MP system (Bio-Rad). Densitometric analysis was performed using Image Lab 6.4 (Bio-Rad) and normalised to loading controls.

### Immunocytochemistry and Confocal imaging

Cells grown on poly-L-lysine-coated coverslips were fixed with 4% paraformaldehyde in PBS for 15 min at room temperature, washed three times with PBS, and permeabilised with 0.2% Triton X-100 in PBS for 10 min. Non-specific binding was blocked with 10% normal goat serum (NGS) or 1% BSA in PBST (0.1% Tween-20) for 1 h. Coverslips were incubated with primary antibodies (diluted in blocking buffer) overnight at 4 °C. The following primary antibodies were used: rabbit anti-phospho-TDP-43 (1:500; Cosmo Bio), mouse anti-CD9 (1:200; BD Biosciences, 555370), early endosomes (rabbit anti-EEA1, 1:500), late endosomes (rabbit anti-Rab7, 1:500), or lysosomes (rabbit anti-LAMP1, 1:500), and rabbit anti-α-tubulin (1:1000; Abcam). After washing, cells were incubated with Alexa Fluor 488-, 555-, or 647-conjugated secondary antibodies (1:500; Invitrogen) for 1 h at room temperature in the dark. Coverslips were mounted using ProLong Diamond Antifade Mountant with DAPI (Invitrogen). Images were acquired using a Zeiss Axiovert 200 M confocal microscope with a 40× or 63× oil immersion objective. Fluorescence intensity was quantified using ImageJ (NIH). For each condition, 10–15 fields of view and at least 50 cells per experiment were analysed across three independent experiments.

To evaluate CD9 membrane trafficking and recycling dynamics, a surface-labelling assay was performed. Live neurons were incubated with anti-CD9 antibody diluted in conditioned medium at 4 °C for 30 min to label surface-exposed CD9. Cells were then washed and shifted to 37 °C to allow internalisation for defined time intervals (15–60 min). Residual surface-bound antibody was removed by brief acid wash (0.2 M acetic acid, 0.5 M NaCl, pH 2.5), followed by fixation. Internalised CD9 was detected using fluorescent secondary antibodies, and the ratio of intracellular to surface CD9 was quantified using fluorescence intensity measurements. This assay enabled assessment of CD9 endocytosis and recycling efficiency under different ESCRT conditions.

### Lysosomal Inhibition Assay

To determine whether CD9 loss was mediated by lysosomal degradation, neurons were treated with Bafilomycin A1 (100 nM) for 4–6 h prior to fixation. Cells were then processed for CD9 immunofluorescence as described above. CD9 fluorescence intensity and puncta characteristics were quantified and compared between treated and untreated conditions. Rescue of CD9 signal following lysosomal inhibition was interpreted as evidence for lysosome-dependent degradation.

### Extracellular Vesicle Isolation

Primary cortical neurons (DIV 7-14) were cultured in Neurobasal medium supplemented with B-27 and GlutaMAX. For EV isolation, cells were washed and incubated in phenol red-free Neurobasal medium with 0.5% B-27 (to reduce serum-derived EV contamination) for 48 h with treatments. Conditioned medium was collected and subjected to differential ultracentrifugation: 300 × g for 10 min to remove cells, 2,000 × g for 20 min to remove debris, and 10,000 × g for 30 min to remove large vesicles. EVs were pelleted at 100,000 × g for 90 min at 4 °C (Optima XE-90 ultracentrifuge, Beckman Coulter). The pellet was washed in PBS and re-centrifuged at 100,000 × g for 90 min. EV pellets were resuspended in RIPA buffer or PBS for downstream analysis.

### Extracellular Vesicle Characterisation

EV preparations were characterised by Western blot. Equal protein amounts (10 µg) of EV lysates were denatured in Laemmli sample buffer and resolved by SDS-PAGE on 12% polyacrylamide gels (Bio-Rad, UK), then transferred onto PVDF membranes (0.45 µm; Bio-Rad) using a wet transfer system. Membranes were blocked in 5% non-fat milk in TBST for 1 h at room temperature, then incubated overnight at 4 °C with primary antibodies against exosomal markers: mouse anti-ALIX (1:500; Santa Cruz), rabbit anti-TSG101 (1:500; Abcam), rabbit anti-VPS4a (1:500; Abcam, ab96488), and the negative control anti-calnexin (1:1000; Abcam, ab22595). After washing with TBST, membranes were incubated with fluorescent secondary antibodies (IRDye 680RD and 800CW; LI-COR) and membranes imaged on a LI-COR Odyssey. Densitometric analysis was performed using Image Studio Lite (LI-COR) and normalised to total protein loaded.

### Dynamic Light Scattering (DLS) Analysis

Dynamic light scattering was used to assess the hydrodynamic diameter and polydispersity of extracellular vesicles generated under each experimental condition. Samples were prepared in sterile phosphate-buffered saline (PBS; pH 7.4, Gibco, UK) and equilibrated at 5 °C for 5 minutes prior to measurement. DLS measurements were performed using a Zetasizer Nano ZS instrument (Malvern Panalytical, UK) equipped with a disposable polystyrene sizing cuvette (Malvern, UK), using a 633 nm He–Ne laser and a fixed scattering angle of 173° (backscatter mode). Each sample was measured for 60 s with automatic attenuator adjustment. The dispersant refractive index (1.330), viscosity (0.8882 cP), and material refractive index (1.45) were applied according to instrument defaults for lipid vesicles. For each experimental condition, six independent biological replicates (n = 6) were analysed. The Z-average diameter (intensity-weighted mean hydrodynamic diameter) and polydispersity index (PdI) were extracted from the cumulants analysis using Zetasizer Software v7.11 (Malvern Panalytical, UK). Samples with PdI > 0.7 were interpreted as highly polydisperse or aggregated. All measurements were performed using the same standard operating procedure (SOP: “DLS liposome 5C.sop”) to ensure consistency across conditions.

## Statistical Analysis

All experiments were performed with at least three independent replicates. Data are presented as mean ± standard deviation (SD) or standard error of the mean (SEM) as indicated in figure legends. Statistical analyses were performed using GraphPad Prism 9 (GraphPad Software, USA). Comparisons between two groups were conducted using unpaired two-tailed Student’s t-test. For comparisons involving three or more groups, one-way or two-way ANOVA was used, followed by appropriate post-hoc tests (Tukey’s or Dunnett’s). Significance was defined as p < 0.05. All data met assumptions of normality and homogeneity of variance; where these were violated, non-parametric alternatives were applied.

## Results

### Alterations in ESCRT Protein Expression in MND Patients

To investigate whether ESCRT components are dysregulated in MND, we examined the expression of key ESCRT proteins in postmortem motor cortex and spinal cord tissues from MND patients and age-matched controls by western blotting. In the motor cortex, CHMP2B, an ESCRT-III subunit involved in intraluminal vesicle formation, was significantly increased by approximately 50% in MND samples compared to controls (p < 0.01; Figure 1A, C). Conversely, TSG101, an ESCRT-I component critical for multivesicular body formation, was significantly decreased in MND motor cortex (p < 0.001; Figure 1A, D). VPS37A expression was also significantly reduced in MND motor cortex (p < 0.05), while VPS37B showed a non-significant trend toward reduction (Figure 1E, F). ALIX and STAM1 expression levels were not significantly altered in the motor cortex (Figure 1A, G).

**Figure 1.**
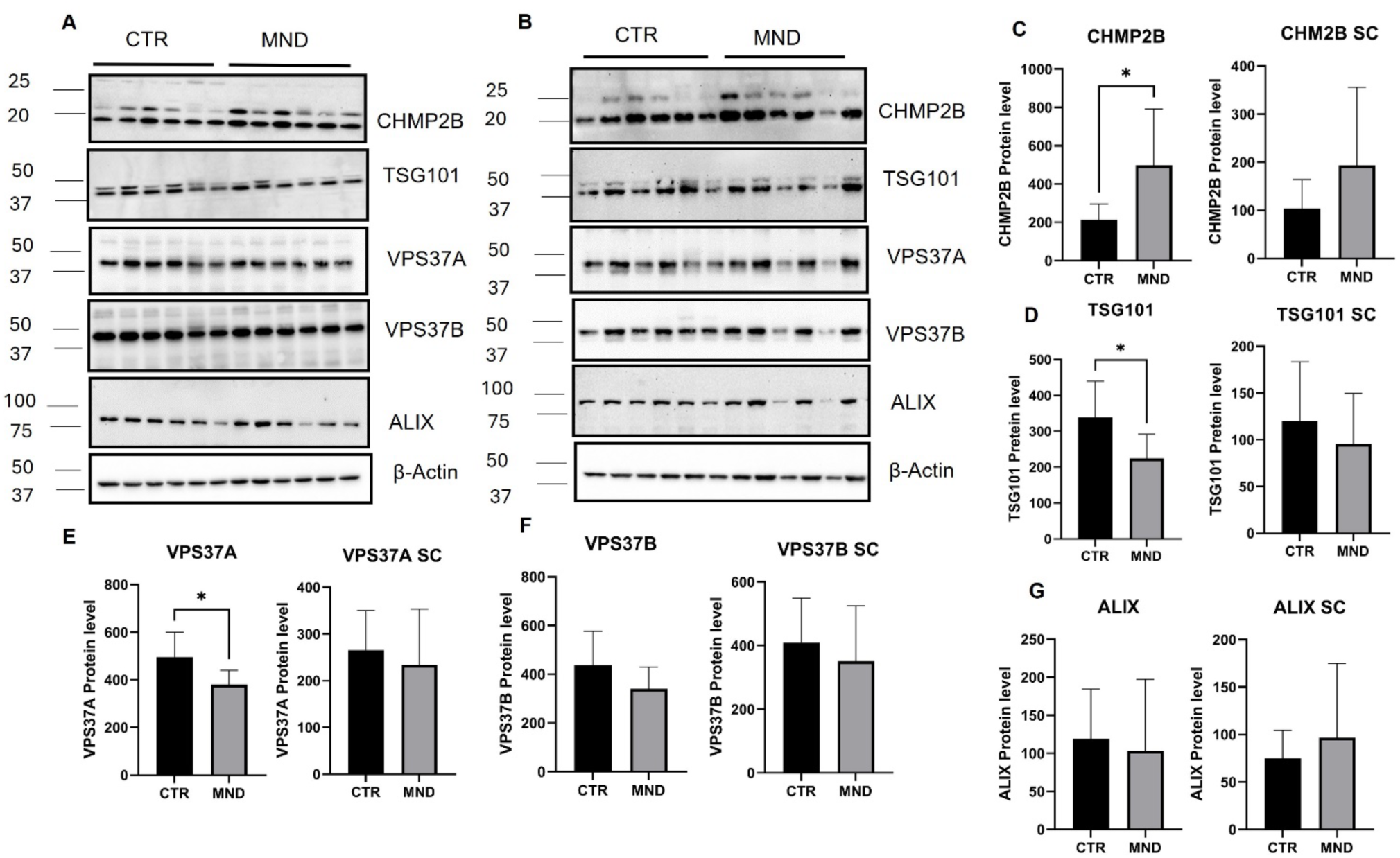
**Expression of ESCRT proteins in MND and control patients**(A) Representative immunoblots of ESCRT proteins (CHMP2B, TSG101, VPS37A, VPS37B, ALIX) in motor cortex lysates from age-matched control (CTR) and MND patients. β-Actin was used as a loading control. (B) Representative immunoblots of ESCRT proteins in spinal cord lysates from control and MND patients. β-Actin served as a loading control. (C) Quantification of CHMP2B protein expression in motor cortex and spinal cord. (D) Quantification of TSG101 protein expression in motor cortex and spinal cord. (E) Quantification of VPS37A protein expression in motor cortex and spinal cord. (F) Quantification of VPS37B protein expression in motor cortex and spinal cord. (G) Quantification of ALIX protein expression in motor cortex and spinal cord.

In the spinal cord (SC), CHMP2B expression was similarly elevated in MND samples compared to controls (p < 0.05; Figure 1B, C). TSG101 expression showed a non-significant trend toward reduction in MND spinal cord (p = 0.08). VPS37A, VPS37B, and ALIX showed no significant changes (Figure 1B, E, F, and G). These findings demonstrate region-specific alterations in ESCRT protein expression in MND, with consistent upregulation of CHMP2B and downregulation of TSG101 in the motor cortex.

Data are presented as mean ± SD. For each region, n = 6 per group (CTR, n = 6; MND, n = 6). Statistical analysis was performed using unpaired two-tailed Student’s t-test. *p < 0.05, p < 0.01, *p < 0.001, ****p < 0.0001. CHMP2B expression was significantly increased in MND motor cortex (p < 0.01) and spinal cord (*p < 0.05) compared to controls. TSG101 expression was significantly decreased in MND motor cortex (*p < 0.001), with a non-significant trend toward reduction in spinal cord. VPS37A expression was significantly reduced in both motor cortex (p < 0.05) and spinal cord (p < 0.05). VPS37B and ALIX showed no significant alterations in either region. SC, spinal cord.

### Tunicamycin increased TDP-43 aggregation in NSC-34 cells

Given that TDP-43 pathology is a hallmark of MND, we first validated our cellular model. Tunicamycin, an ER stress inducer, has been shown to promote TDP-43 accumulation. Treatment of differentiated NSC-34 cells with 100 nM tunicamycin for 48 h significantly increased both total TDP-43 (tTDP-43) and phosphorylated TDP-43 (pTDP-43 Ser409/410) levels compared to vehicle-treated controls (tTDP-43: p < 0.01; pTDP-43: p < 0.001; Figure 2B1–B3). Immunocytochemistry confirmed increased pTDP-43 immunoreactivity in tunicamycin-treated cells (p < 0.01; Figure 2C1–C3). Consistent with human MND pathology, postmortem MND brain tissues exhibited elevated pTDP-43 levels compared to controls (p < 0.01; Figure 2A1, A2), confirming the relevance of this model.

**Figure 2.**
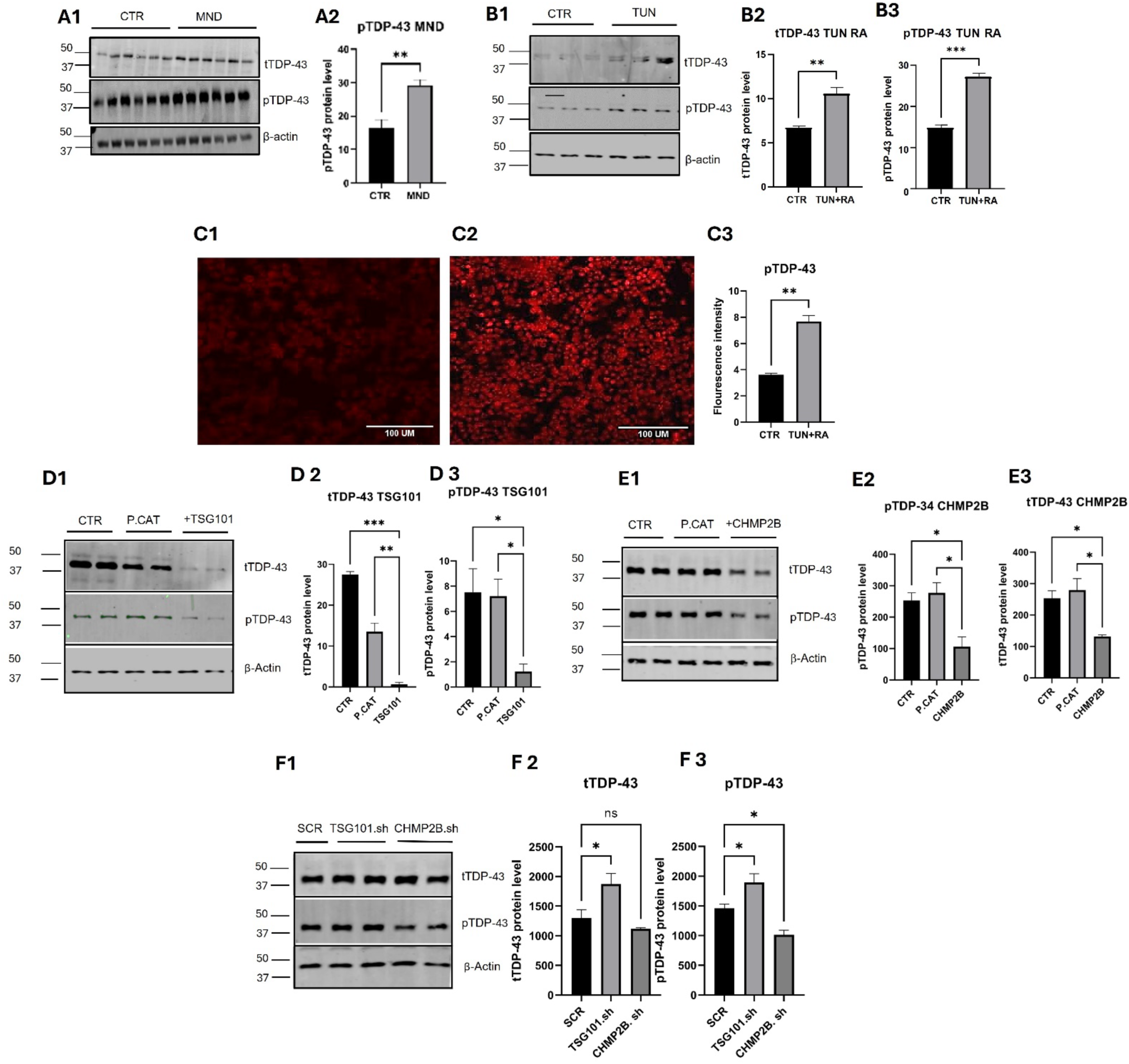
Tunicamycin increases TDP-43 aggregation in NSC-34 cells and TSG101/CHMP2B differentially regulate TDP-43 accumulation and phosphorylation. (A1) Representative immunoblots of total TDP-43 (tTDP-43) and phosphorylated TDP-43 (pTDP-43 Ser409/410) in motor cortex lysates from control (CTR) and MND patients. β-Actin is the loading control. (A2) Quantification of pTDP-43 in control vs. MND brains (n = 6 per group). pTDP-43 was significantly increased in MND brains (p = 0.0017, unpaired two-tailed t-test). (B1) Representative immunoblots of tTDP-43 and pTDP-43 in differentiated NSC-34 cells treated with vehicle (CTR) or 100 nM tunicamycin (TUN) for 48 h in the presence of retinoic acid (RA). β-Actin is the loading control. (B2) Quantification of tTDP-43. TUN significantly increased tTDP-43 compared to CTR (p = 0.006). (B3) Quantification of pTDP-43. TUN significantly increased pTDP-43 compared to CTR (p = 0.0003). (C1) Immunocytochemistry of pTDP-43 (red) in control differentiated NSC-34 cells. Scale bar = 100 µm. (C2) Immunocytochemistry of pTDP-43 (red) in NSC-34 cells treated with TUN (100 nM, 48 h) + RA.(C3) Quantification of pTDP-43 fluorescence intensity. TUN significantly increased pTDP-43 immunoreactivity compared to CTR (p = 0.0065). (D1) Representative immunoblots of tTDP-43 and pTDP-43 in differentiated NSC-34 cells exposed to TUN (100 nM, 48 h) under the following conditions: CTR (tunicamycin-treated), P.CAT (empty plasmid control), and TSG101 (TSG101-overexpressing). β-Actin is the loading control. (D2) Quantification of tTDP-43. TSG101 overexpression significantly reduced tTDP-43 compared to P.CAT (p = 0.0318). (D3) Quantification of pTDP-43. TSG101 overexpression significantly reduced pTDP-43 compared to P.CAT (p = 0.0318). (E1) Representative immunoblots of tTDP-43 and pTDP-43 in differentiated NSC-34 cells exposed to TUN (100 nM, 48 h) under the following conditions: CTR, P.CAT, and CHMP2B (CHMP2B-overexpressing). β-Actin is the loading control. (E2) Quantification of tTDP-43. CHMP2B overexpression significantly reduced tTDP-43 compared to P.CAT (p = 0.0189). (E3) Quantification of pTDP-43. CHMP2B overexpression significantly reduced pTDP-43 compared to P.CAT (p = 0.0188). (F1) Representative immunoblots of tTDP-43 and pTDP-43 in differentiated NSC-34 cells transfected with scrambled shRNA (SCR), TSG101 shRNA, or CHMP2B shRNA under TUN-induced ER stress. β-Actin is the loading control. (F2) Quantification of tTDP-43. TSG101 knockdown significantly increased tTDP-43 compared to SCR (p = 0.0188); CHMP2B knockdown had no significant effect. (F3) Quantification of pTDP-43. CHMP2B knockdown significantly reduced pTDP-43 compared to SCR (p = 0.0029); TSG101 knockdown had no significant effect. All data are mean ± SD from 3 independent experiments. Two-group comparisons (A2, B2, B3, C3) used unpaired two-tailed Student’s t-test. Multi-group comparisons (D2, D3, E2, E3, F2, F3) used one-way ANOVA with appropriate post-hoc test. Significance: *p < 0.05, **p < 0.01, ***p < 0.001, ****p < 0.0001. Abbreviations: CTR, control; TUN, tunicamycin; RA, retinoic acid; tTDP-43, total TDP-43; pTDP-43, phosphorylated TDP-43 (Ser409/410); P.CAT, empty plasmid control; SCR, scrambled shRNA control.

### TSG101 and CHMP2B Differentially Regulate TDP-43 Accumulation and Phosphorylation

To determine whether modulating ESCRT components influences TDP-43 pathology, we overexpressed TSG101 or CHMP2B in differentiated NSC-34 cells under tunicamycin-induced ER stress. Overexpression of TSG101 significantly reduced both tTDP-43 (p < 0.05) and pTDP-43 (p < 0.05) levels compared to control plasmid-transfected cells (Figure 2D1–D3). Similarly, CHMP2B overexpression significantly decreased tTDP-43 (p < 0.05) and pTDP-43 (p < 0.05) levels (Figure 2E1–E3). Conversely, TSG101 knockdown using shRNA significantly increased tTDP-43 levels (p < 0.05) compared to the scrambled control (Figure 2F1, F2), indicating that TSG101 is required for limiting TDP-43 accumulation. In contrast, CHMP2B knockdown significantly reduced pTDP-43 levels (p < 0.01) without significantly altering tTDP-43 (Figure 2F1, F3), suggesting that CHMP2B may specifically influence TDP-43 phosphorylation rather than total protein levels. Collectively, these data indicate that increasing the expression of specific ESCRT components can mitigate TDP-43 pathology under proteostatic stress, while TSG101 and CHMP2B exert distinct regulatory effects on TDP-43 accumulation and phosphorylation.

### ESCRT Proteins Modulate mTOR Signaling and Autophagy Markers

To explore the mechanism by which ESCRT components regulate TDP-43, we examined the effects of TSG101 and CHMP2B modulation on the mTOR pathway and autophagy markers. Overexpression of TSG101 significantly reduced both total mTOR (t-mTOR) (p < 0.001) and phosphorylated mTOR (p-mTOR) (p < 0.01) levels (Figure 3A1–A3). Conversely, TSG101 knockdown increased p-mTOR levels (p < 0.01). CHMP2B overexpression also reduced p-mTOR levels (p < 0.01), while CHMP2B knockdown increased p-mTOR (p < 0.05) (Figure 3A1–A3). Regarding autophagy markers, TSG101 overexpression reduced p62 levels (p < 0.001), while TSG101 knockdown increased p62 (Figure 3B1–B3). LC3B-II levels were significantly increased upon TSG101 knockdown (p < 0.01), consistent with autophagic flux impairment (Figure 3C1–C4). CHMP2B modulation showed similar trends. These results indicate that ESCRT proteins regulate TDP-43 at least in part through modulation of mTOR-dependent autophagy.

**Figure 3.**
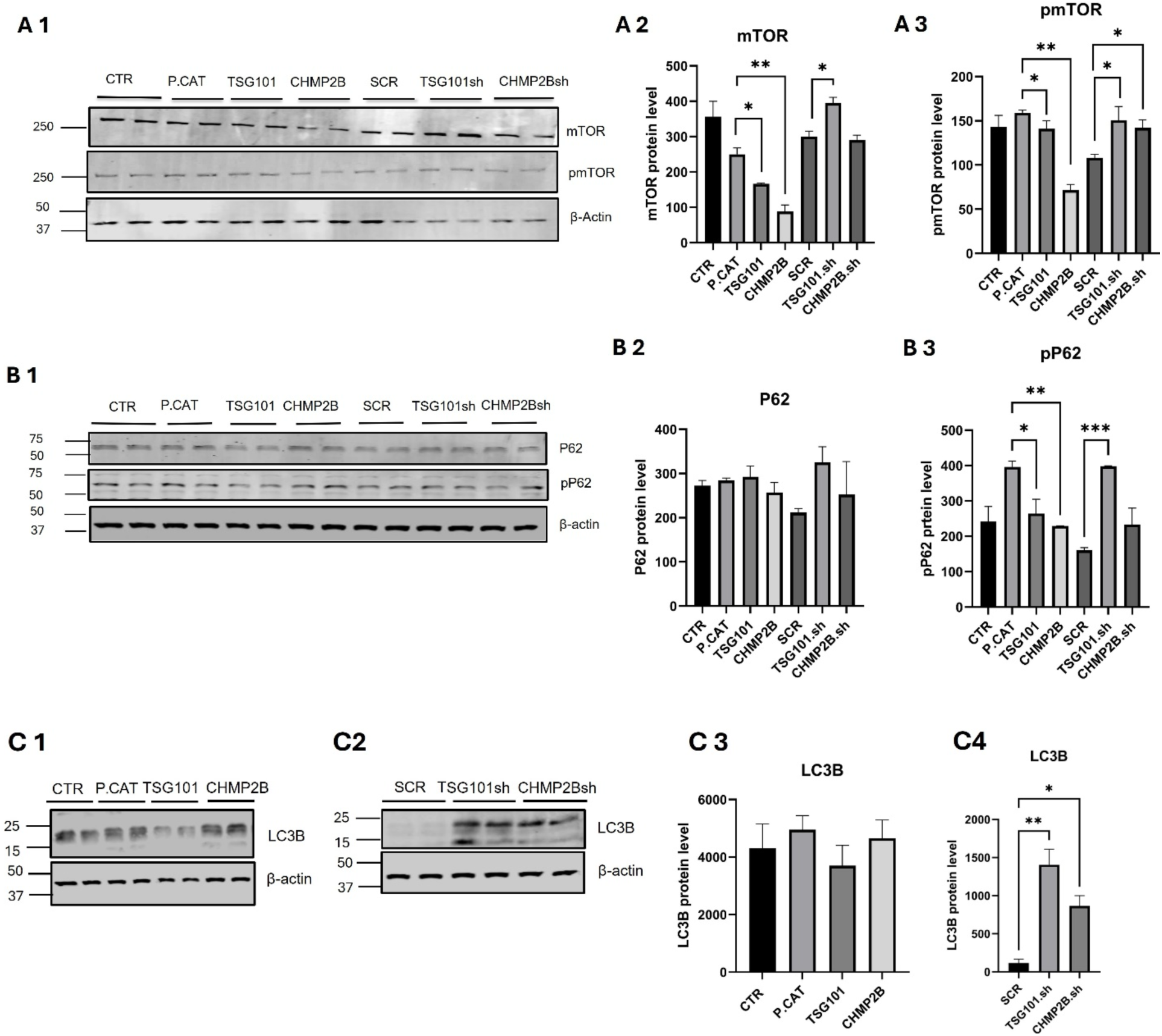
Effect of ESCRT proteins on mTOR, p62, and LC3B expression in NSC-34 cells. (A) TSG101 and CHMP2B modulate mTOR expression.(A1) Representative immunoblots of total mTOR (tmTOR) and phosphorylated mTOR (p-mTOR Ser2448) in differentiated NSC-34 cells under the following conditions: CTR (control, tunicamycin-treated), P.CAT (control plasmid-transfected), TSG101 (TSG101-overexpressing), CHMP2B (CHMP2B-overexpressing), SCR (scrambled shRNA control), TSG101 shRNA (TSG101 knockdown), and CHMP2B shRNA (CHMP2B knockdown). β-Actin was used as a loading control. (A2) Quantification of tmTOR levels normalised to β-actin. Significant differences were observed across conditions (p = 0.0002). (A3) Quantification of p-mTOR levels normalised to β-actin. Significant differences were observed across conditions (p = 0.0036). (B) ESCRT modulation alters p62 expression. (B1) Representative immunoblots of total p62 (tp62) and phosphorylated p62 (pp62) in NSC-34 cells under the indicated conditions. β-Actin served as a loading control. (B2) Quantification of tp62 levels normalised to β-actin. No significant differences were observed (p = 0.1650). (B3) Quantification of pp62 levels normalised to β-actin. Significant differences were observed across conditions (p = 0.0005). (C) LC3B expression is differentially regulated by TSG101 and CHMP2B. (C1) Representative immunoblots of LC3B in NSC-34 cells under overexpression conditions (CTR, P.CAT, TSG101, CHMP2B). β-Actin was used as a loading control. (C2) Representative immunoblots of LC3B in NSC-34 cells under knockdown conditions (SCR, TSG101 shRNA, CHMP2B shRNA). β-Actin served as a loading control. (C3) Quantification of LC3B levels normalised to β-actin under overexpression conditions. No significant differences were observed (p = 0.4073). (C4) Quantification of LC3B levels normalised to β-actin under knockdown conditions. TSG101 knockdown significantly increased LC3B levels compared to SCR (p = 0.0070). All data are presented as mean ± SD from at least three independent experiments. CTR, control (tunicamycin-treated + retinoic acid); P.CAT, empty plasmid control; TSG101, TSG101-overexpressing; CHMP2B, CHMP2B-overexpressing; SCR, scrambled shRNA control; TSG101 shRNA, TSG101 knockdown; CHMP2B shRNA, CHMP2B knockdown. Statistical significance was determined using one-way ANOVA with appropriate post-hoc test. *p < 0.05, **p < 0.01, ***p < 0.001, ****p < 0.0001.

### Downregulation of TSG101 increases TDP-43 in primary neuronal cells

To validate our findings in a more physiologically relevant system, we transduced primary cortical neurons with lentivirus expressing TSG101 shRNA or control shRNA. TSG101 knockdown significantly increased both tTDP-43 (p < 0.01) and pTDP-43 (p < 0.01) levels compared to controls (Figure 4A, B1, B2). This confirms that TSG101 deficiency promotes TDP-43 accumulation in primary neurons.

**Figure 4.**
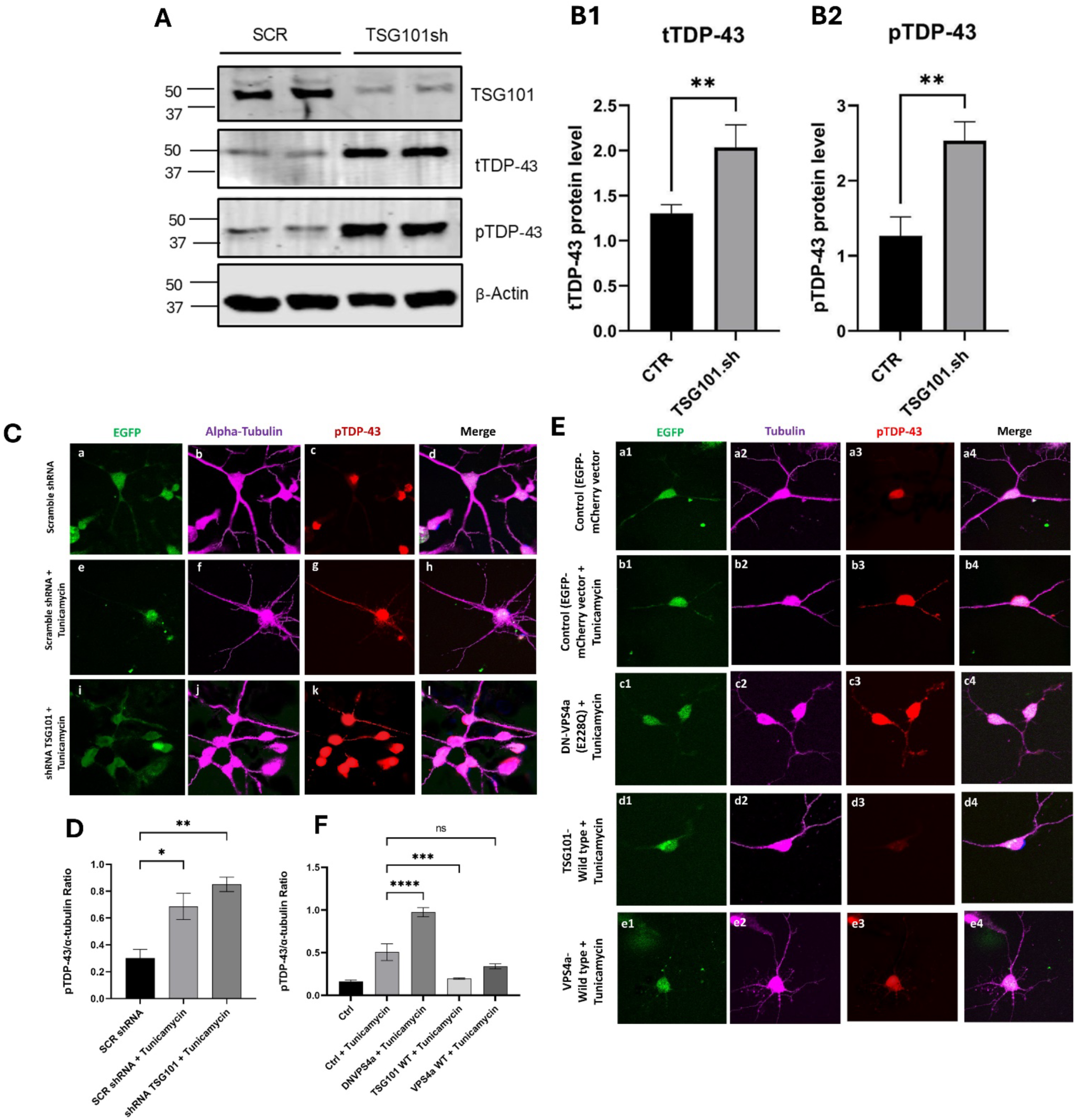
Downregulation of TSG101 increases TDP-43 expression and ESCRT modulation alters pTDP-43 localisation under ER stress in primary neurons. (A) Representative immunoblots of total TDP-43 (tTDP-43) and phosphorylated TDP-43 (pTDP-43 Ser409/410) in primary cortical neurons infected with control lentivirus (CTR) or TSG101 shRNA lentivirus (TSG101.sh). β-Actin was used as a loading control. (B1) Quantification of tTDP-43 levels normalised to β-actin. TSG101 knockdown significantly increased tTDP-43 expression compared to control (p = 0.0094). (B2) Quantification of pTDP-43 levels normalised to β-actin. TSG101 knockdown significantly increased pTDP-43 expression compared to control (p = 0.0035). All data are presented as mean ± SD from at least three independent experiments. CTR, primary neurons infected with control lentivirus; TSG101.sh, primary neurons infected with TSG101 shRNA lentivirus. Statistical significance was determined using unpaired two-tailed Student’s t-test. *p < 0.05, **p < 0.01, ***p < 0.001, ****p < 0.0001. (C) ESCRT depletion enhances pTDP-43 mislocalisation. Confocal images of primary hippocampal neurons labeled for EGFP (green), α-tubulin (magenta), and pTDP-43 (red). a–d, untreated controls showing predominantly nuclear pTDP-43 localisation. e–h, tunicamycin-treated neurons (100 nM, 48 h) exhibiting mild cytoplasmic pTDP-43 redistribution. i–l, neurons expressing TSG101 shRNA (shTSG101) combined with tunicamycin treatment, showing pronounced cytoplasmic accumulation of pTDP-43. Scale bar = 10 µm. (D) Quantification of pTDP-43 fluorescence intensity normalised to α-tubulin. Data are presented as mean ± SEM from n = 6 cells per condition across three independent experiments. Statistical analysis was performed using one-way ANOVA with Tukey’s post-hoc test. *p < 0.05, **p < 0.01. (E) Modulation of ESCRT components alters pTDP-43 localisation. Confocal images of neurons labeled for EGFP (green), α-tubulin (magenta), and pTDP-43 (red/blue). a1–a4, untreated controls with nuclear pTDP-43 localisation. b1–b4, control (EGFP–mCherry) neurons treated with tunicamycin, showing partial cytoplasmic redistribution. c1–c4, neurons expressing dominant-negative VPS4a (DN-VPS4a, E228Q) combined with tunicamycin, exhibiting marked cytoplasmic pTDP-43 mislocalisation. d1–d4, neurons overexpressing wild-type TSG101 (TSG101-WT) combined with tunicamycin, showing reduced cytoplasmic pTDP-43 and preserved nuclear localisation. e1–e4, neurons overexpressing wild-type VPS4a (VPS4a-WT) combined with tunicamycin, also limiting cytoplasmic accumulation. Scale bar = 10 µm. (F) Quantification of pTDP-43/α-tubulin fluorescence intensity ratio. Data are presented as mean ± SEM from n = 3 independent experiments. Statistical analysis was performed using one-way ANOVA with Tukey’s post-hoc test. ****p < 0.0001, ***p < 0.001 compared to control + tunicamycin condition; ns, not significant.

### Modulation of ESCRT Components Influences TDP-43 Localisation under ER Stress

To examine whether ESCRT dysfunction alters TDP-43 pathology under ER stress, the effects of TSG101 knockdown on phosphorylated TDP-43 (pTDP-43) accumulation were tested in primary cortical neurons. Under basal conditions, neurons displayed diffuse nuclear pTDP-43 immunoreactivity (Figure 4C(a–d)). Following tunicamycin treatment, control neurons showed increased cytoplasmic redistribution of pTDP-43, accompanied by loss of nuclear staining (Fig. 4C(e–h)). This effect was markedly enhanced in shTSG101-expressing neurons, which exhibited robust cytoplasmic pTDP-43 inclusions and reduced nuclear retention (Fig. 4C(i–l)). Quantification confirmed a significant increase in cytoplasmic pTDP-43 fluorescence intensity normalised to α-tubulin in TSG101-depleted neurons (p < 0.01; Figure 4D). These data indicate that ESCRT-I deficiency potentiates stress-induced mislocalisation of TDP-43.

To further investigate the role of ESCRT machinery in TDP-43 localisation, we expressed dominant-negative VPS4a (DN-VPS4a, E228Q), wild-type TSG101, or wild-type VPS4a in primary neurons. DN-VPS4a expression under tunicamycin treatment resulted in pronounced cytoplasmic accumulation of pTDP-43 compared to vector controls (Figure 4E(c1–c4) vs. (b1–b4)). Quantification revealed that DN-VPS4a significantly increased the pTDP-43/α-tubulin ratio (p < 0.0001; Figure 4F). In contrast, overexpression of TSG101-WT or VPS4a-WT partially rescued tunicamycin-induced cytoplasmic pTDP-43 aggregation, preserving nuclear localisation (Figure 4E(d1–d4), (e1–e4)). TSG101-WT overexpression resulted in the most pronounced protection, with pTDP-43/α-tubulin ratios significantly lower than control-treated cells (p < 0.0001; Figure 4F). These findings suggest that ESCRT components play protective roles in limiting pathological TDP-43 mislocalisation under proteostatic stress.

### ESCRT Proteins Differentially Regulate CD9 Tetraspanin Levels During ER Stress

To investigate how ESCRT machinery components influence membrane protein stability during cellular stress, the effects of TSG101 and VPS4a manipulation on CD9 tetraspanin expression were examined under basal and ER stress conditions in primary neurons. CD9 is a tetraspanin protein involved in membrane organisation and cellular signaling, making it a relevant target for studying ESCRT-mediated protein regulation. The subcellular localisation of CD9 was examined in relation to early endosomes (EEA1), late endosomes (Rab7), and lysosomes (LAMP1) under basal and ER stress conditions in primary cultured neurons (Figure 5).

**Figure 5.**
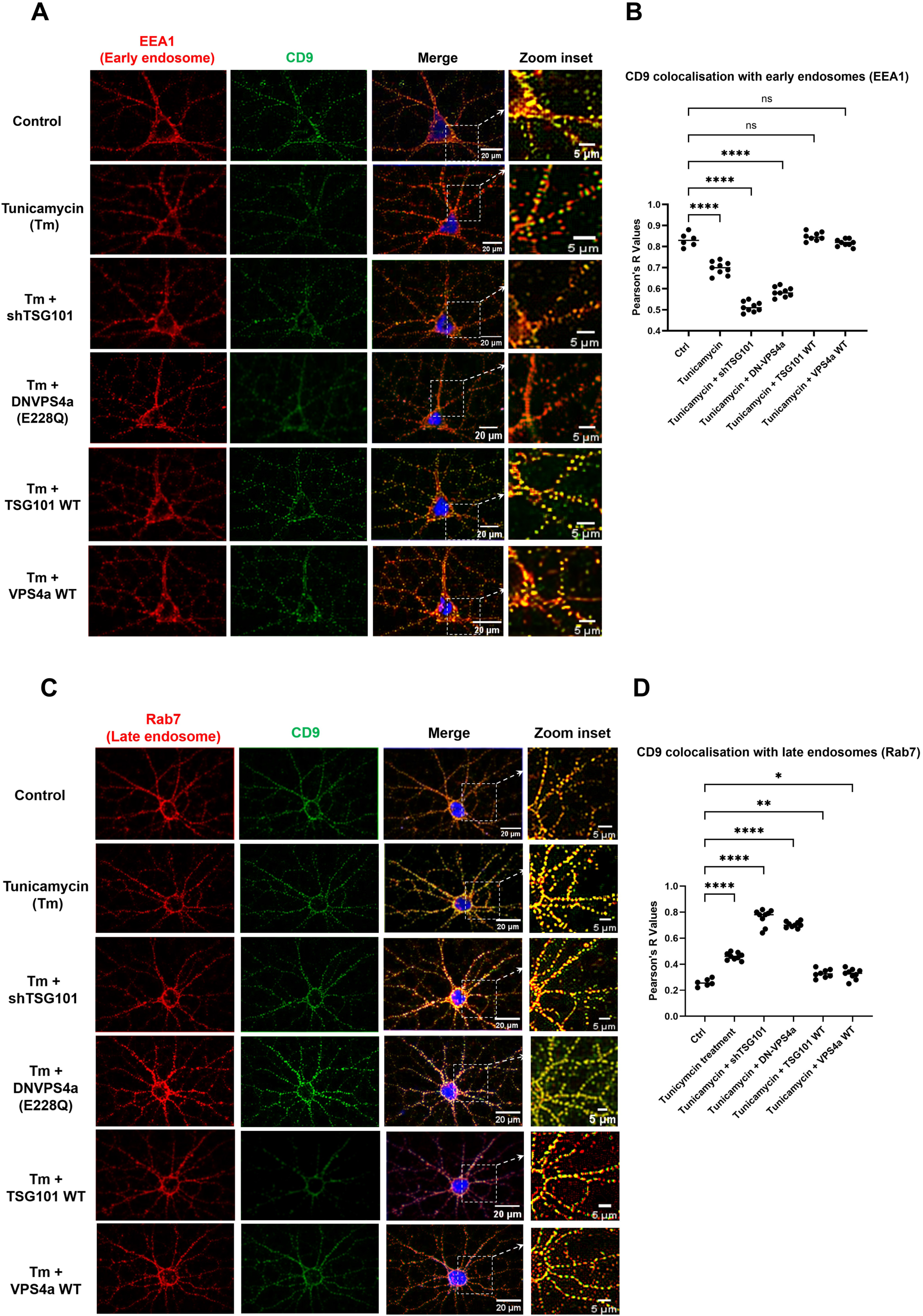
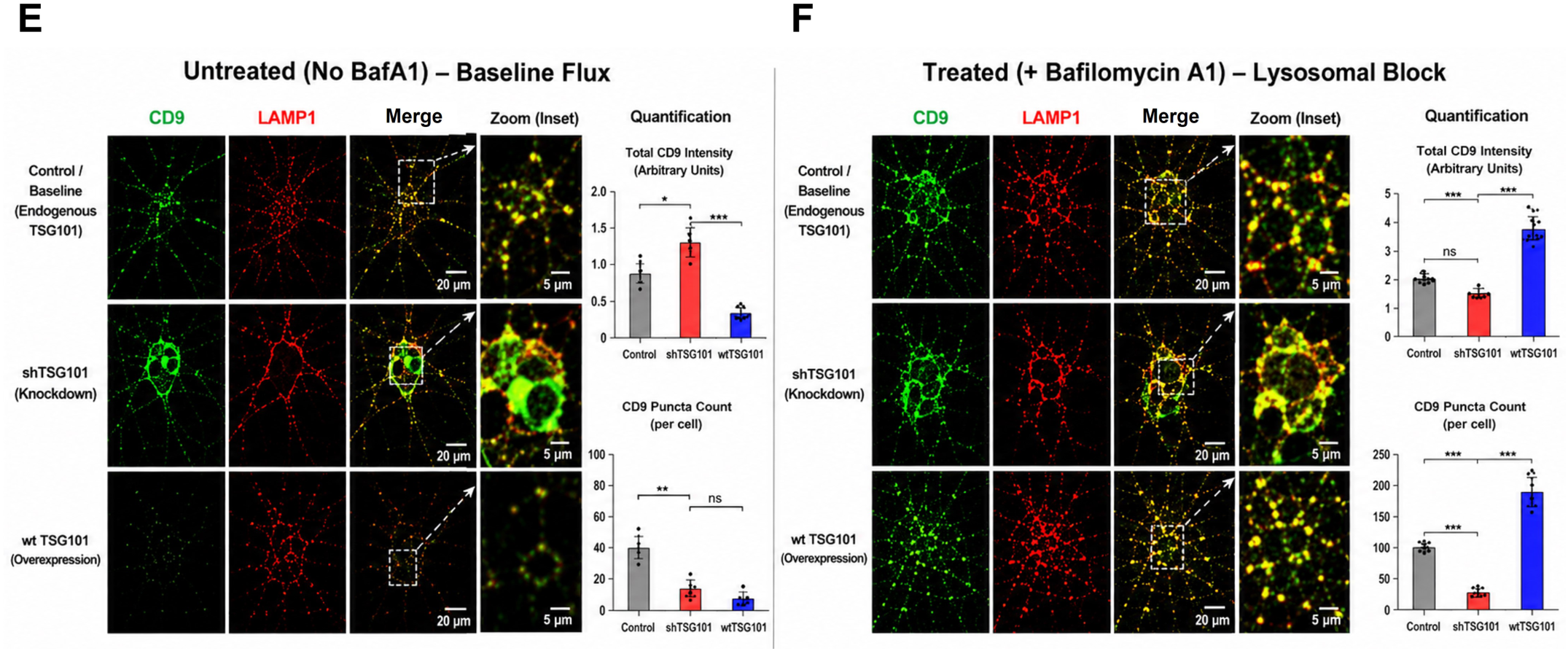
ESCRT modulation alter CD9-positive vesicle trafficking linked to EV biogenesis in cortical primary neurons under ER stress. (A) Representative confocal immunofluorescence images showing the colocalisation of CD9 (green) with the early endosomal marker EEA1 (red) in primary cortical neurons under control conditions or following tunicamycin (Tm) treatment in combination with modulation of ESCRT function (shTSG101, dominant-negative VPS4a [DNVPS4a, E228Q], TSG101 WT, or VPS4a WT). Merged images demonstrate areas of colocalisation (yellow/orange), while boxed regions are shown at higher magnification in the corresponding zoom insets. Under basal conditions, CD9 exhibits substantial localisation within early endosomes. Tunicamycin treatment significantly reduces CD9 accumulation in EEA1-positive compartments, whereas disruption of ESCRT function by shTSG101 or DNVPS4a further decreases early endosomal localisation. In contrast, overexpression of WT TSG101 or VPS4a restores CD9 trafficking to early endosomes. (B) Quantification of CD9 colocalisation with EEA1 expressed as Pearson’s correlation coefficient. Tunicamycin significantly decreases CD9–EEA1 colocalisation compared with untreated controls. ESCRT inhibition (shTSG101 or DNVPS4a) further reduces colocalisation, whereas WT TSG101 or VPS4a significantly restores early endosomal localisation toward control levels. Each point represents an individual cell; horizontal bars indicate mean ± SEM. Statistical analysis was performed using one-way ANOVA followed by Tukey’s multiple-comparison test. ns, not significant; ****P < 0.0001. (C) Representative confocal immunofluorescence images showing the colocalisation of CD9 (green) with the late endosomal marker Rab7 (red) under the same experimental conditions. Tunicamycin promotes redistribution of CD9 from early to late endosomes, resulting in increased CD9–Rab7 colocalisation. Impairment of ESCRT function by shTSG101 or DNVPS4a further enhances CD9 accumulation within Rab7-positive compartments, whereas WT TSG101 or VPS4a overexpression reduces late endosomal retention by promoting efficient trafficking toward lysosomal degradation. (D) Quantification of CD9 colocalisation with Rab7 expressed as Pearson’s correlation coefficient. Tunicamycin significantly increases CD9 localisation within late endosomes, with a further increase following ESCRT inhibition. Restoration of ESCRT activity by WT TSG101 or VPS4a significantly decreases CD9–Rab7 colocalisation, consistent with enhanced progression through the endosomal maturation pathway. Data are presented as mean ± SEM. Statistical significance was determined by one-way ANOVA with Tukey’s post hoc test (*P < 0.05, **P < 0.01, ****P < 0.0001). (E) Immunofluorescence images illustrating CD9 (green) and the lysosomal marker LAMP1 (red) in tunicamycin-treated neurons under basal lysosomal flux (without Bafilomycin A1). Control cells exhibit moderate CD9 fluorescence with a moderate number of lysosome-associated puncta, reflecting constitutive lysosomal degradation. TSG101 knockdown impairs lysosomal targeting, resulting in enlarged CD9-positive aggregates with reduced puncta numbers and limited CD9–LAMP1 colocalisation. Conversely, WT TSG101 overexpression is markedly reduce CD9 fluorescence because of accelerated ESCRT-dependent lysosomal degradation. Bar graphs illustrate the total CD9 fluorescence intensity and puncta number per cell. (F) Immunofluorescence images illustrating CD9 and LAMP1 following treatment with Bafilomycin A1, which inhibits lysosomal acidification and degradation. In control neurons, Bafilomycin A1 causes intracellular accumulation of CD9-positive lysosomal puncta with increased CD9–LAMP1 colocalisation. Cells expressing shTSG101 exhibit minimal additional accumulation because defective ESCRT function prevents efficient lysosomal delivery, leading to persistent CD9 aggregates. In contrast, WT TSG101 overexpression produces the greatest increase in CD9 fluorescence and puncta following lysosomal blockade, reflecting efficient lysosomal targeting coupled with inhibition of degradation. Bar graphs summarize the changes in total CD9 fluorescence intensity and puncta number. Scale bars: 20 μm (main images) and 5 μm (magnified insets). Abbreviations: Tm, tunicamycin; TSG101, Tumor Susceptibility Gene 101; VPS4a, Vacuolar Protein Sorting-associated Protein 4A; DNVPS4a, dominant-negative VPS4a (E228Q); EEA1, Early Endosome Antigen 1; Rab7, Ras-related protein Rab7; LAMP1, Lysosome-associated membrane protein 1; BafA1, Bafilomycin A1; CD9, Cluster of Differentiation 9.

Under control conditions, CD9 showed strong colocalisation with the early endosome marker EEA1, with a Pearson’s correlation coefficient of 0.83 ± 0.03 (Figure 5A, B). Tunicamycin treatment significantly reduced CD9-EEA1 colocalisation to 0.70 ± 0.03 (p < 0.001 vs. control), indicating that ER stress disrupts the normal association of CD9 with early endosomes. TSG101 knockdown (shTSG101) and DN-VPS4a expression further reduced colocalisation to 0.51 ± 0.02 and 0.58 ± 0.02, respectively (p < 0.001 vs. tunicamycin). In contrast, both TSG101 WT and VPS4a WT overexpression restored CD9-EEA1 colocalisation to control levels (TSG101 WT: 0.84 ± 0.02; VPS4a WT: 0.82 ± 0.02; ns vs. control for both). These data demonstrate that restoring either ESCRT-I (TSG101) or ESCRT-III (VPS4a) function protects CD9 localisation to early endosomes under ER stress.

Tunicamycin treatment significantly increased CD9 colocalisation with the late endosome marker Rab7 from 0.26 ± 0.03 in control neurons to 0.46 ± 0.03 (p < 0.001 vs. control; Figure 5C, D). TSG101 knockdown and DN-VPS4a expression further increased CD9-Rab7 colocalisation to 0.78 ± 0.02 and 0.70 ± 0.02, respectively (p < 0.001 vs. tunicamycin), indicating pronounced accumulation of CD9 in late endosomal compartments when ESCRT function is impaired. Notably, both TSG101 WT and VPS4a WT overexpression reduced CD9-Rab7 colocalisation to control levels (TSG101 WT: 0.33 ± 0.03; VPS4a WT: 0.34 ± 0.03; ns vs. control for both), suggesting that restoring either ESCRT-I or ESCRT-III function rescues CD9 trafficking through the late endosomal pathway.

Under control conditions, minimal CD9 colocalisation with the lysosomal marker LAMP1 was observed (Pearson’s R = 0.13 ± 0.02; Figure 5E, F). Tunicamycin treatment significantly increased CD9-LAMP1 colocalisation to 0.36 ± 0.03 (p < 0.01 vs. control), indicating enhanced lysosomal targeting of CD9 under ER stress. TSG101 knockdown markedly increased CD9-LAMP1 colocalisation to 0.65 ± 0.02 (p < 0.0001 vs. tunicamycin), while DN-VPS4a expression produced a moderate increase to 0.42 ± 0.02 (p < 0.01 vs. tunicamycin). Strikingly, both TSG101 WT and VPS4a WT overexpression significantly reduced CD9-LAMP1 colocalisation to control levels (TSG101 WT: 0.19 ± 0.02; VPS4a WT: 0.19 ± 0.02; p < 0.001 vs. tunicamycin; ns vs. control for both).

Collectively, these results demonstrate that ER stress disrupts normal CD9 trafficking, leading to reduced association with early endosomes and increased accumulation in late endosomes and lysosomes. TSG101 knockdown and DN-VPS4a expression exacerbates this phenotype, trapping CD9 in degradative compartments. Importantly, both TSG101 WT and VPS4a WT overexpression rescue CD9 trafficking, restoring early endosome association and reducing late endosomal and lysosomal targeting to control levels. These findings establish that proper ESCRT function that is mediated by both ESCRT-I (TSG101) and ESCRT-III (VPS4a) is essential for maintaining CD9 trafficking and preventing its stress-induced mislocalisation to degradative pathways.

### Distinct ESCRT Pathways Regulate Exosome Biogenesis and Pathological TDP-43 Secretion in MND Model

#### ESCRT-dependent loading of TDP-43 into extracellular vesicles

To investigate the role of the ESCRT pathway in TDP-43 secretion, the purity of extracellular vesicle (EV) preparations was first characterised. Immunoblot analysis confirmed the enrichment of canonical exosomal markers, including ALIX, TSG101, and VPS4a, in the EV fraction. The absence of the endoplasmic reticulum protein Calnexin indicated that the preparations were free of significant contaminating organelles (Figure 6).

**Figure 6.**
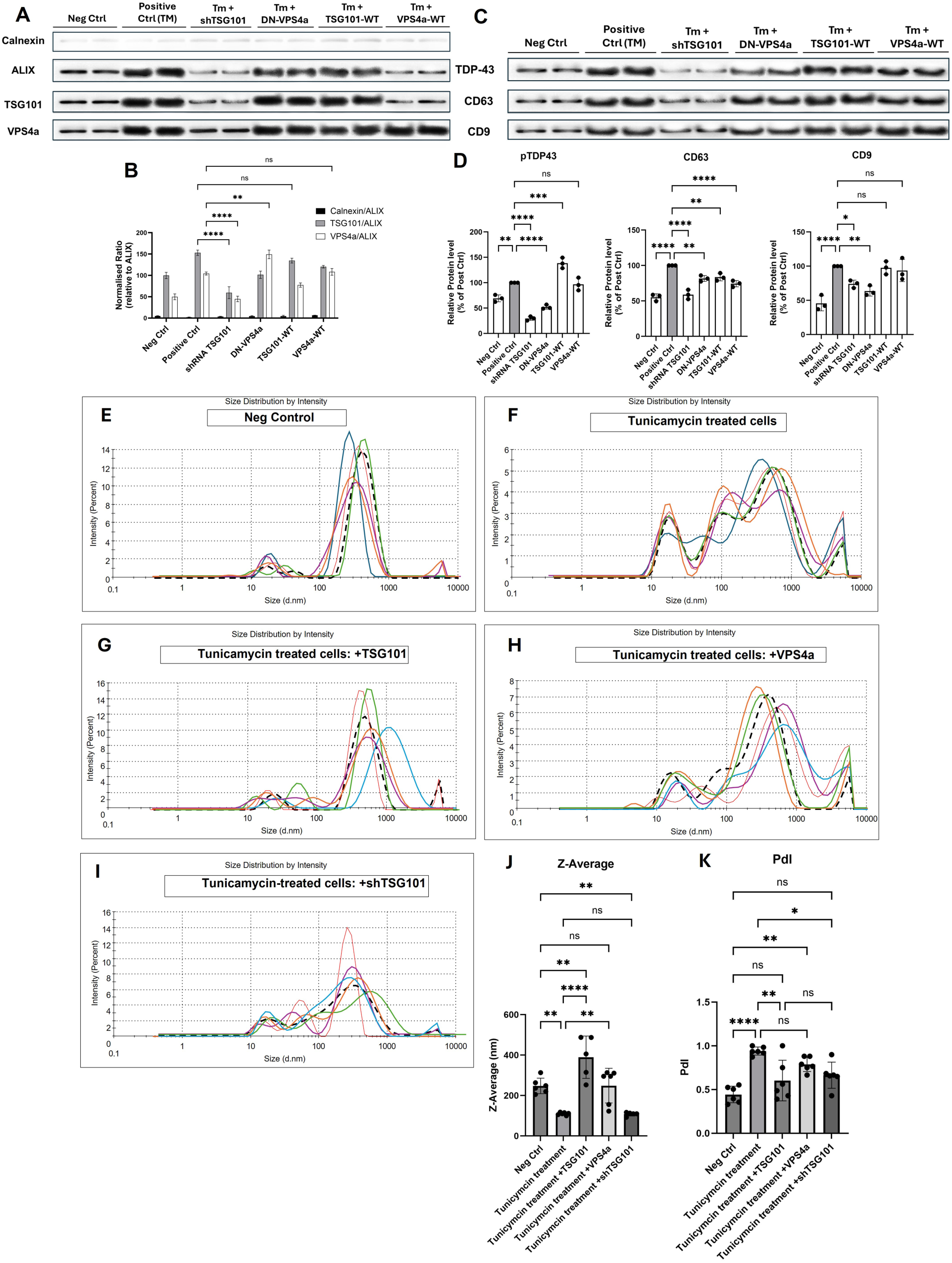
ESCRT components differentially regulate extracellular vesicles biogenesis, pathological TDP-43 loading, and physical properties of vesicles in primary neurons under ER stress. (A) Validation of extracellular vesicle (EV) isolation purity. Blot analysis of exosome-enriched EVs isolated from culture media of tunicamycin-treated primary neurons under various experimental conditions. EVs were characterised using established markers: ALIX, TSG101 (ESCRT-associated proteins), and VPS4a (AAA-ATPase involved in ESCRT disassembly). Calnexin (endoplasmic reticulum marker) was used as a negative control to assess preparation purity. Experimental conditions: negative control (untreated), positive control (tunicamycin 100 nM, 48 h), TSG101 knockdown (shRNA-TSG101), dominant-negative VPS4a (DN-VPS4a, E228Q), wild-type TSG101 overexpression (TSG101-WT), and wild-type VPS4a overexpression (VPS4a-WT). The absence of calnexin signal confirms successful EV isolation with minimal cellular contamination. Data are representative of three independent experiments. (B) Densitometric quantification of blot signals from panel A. Relative protein levels for Calnexin, TSG101, and VPS4a across all experimental conditions. Statistical analysis was performed using ordinary two-way ANOVA with Dunnett’s post-hoc test compared to positive control. **p < 0.01, ****p < 0.0001. (C) ESCRT modulation alters pTDP-43, CD63, and CD9 levels in EVs. Blot showing phosphorylated TDP-43 (pTDP-43), CD63, and CD9 expression across experimental conditions (as in A, B). CD63 and CD9 serve as exosome markers, while pTDP-43 represents MND-associated pathological protein cargo. (D) Summary bar graphs comparing relative protein levels of pTDP-43, CD63, and CD9 across all experimental conditions. Data were normalised to the positive control condition (set to 100%). Values represent mean ± SEM (n = 3 per condition). Statistical analysis was performed using ordinary two-way ANOVA with Dunnett’s post-hoc test compared to positive control. *p <0.05, **p < 0.01, ***p < 0.001, ****p < 0.0001. (E-I) Dynamic light scattering (DLS) analysis of EV size and polydispersity. Representative DLS intensity-based size distribution profiles of EVs isolated from primary neuron culture media under selected conditions: negative control (untreated); positive control (tunicamycin); TSG101 overexpression (+TSG101); TSG101 knockdown (shTSG101); VPS4a overexpression (+VPS4a). Profiles show particle size (d.nm) on the x-axis (log scale) and intensity (%) on the y-axis. Negative control samples displayed a relatively monodisperse peak centred at ∼250 nm (polydispersity index, PdI = 0.44 ± 0.09). Positive control (tunicamycin) generated markedly smaller particles (∼112 nm) with a broad, multi-modal distribution (PdI = 0.94 ± 0.04), indicating a highly heterogeneous mixture. TSG101 overexpression produced the largest vesicles (Z-average: 389 nm) with a broad peak (PdI up to 1.00). TSG101 knockdown resulted in predominantly small vesicles (Z-average: 109 nm) with moderate PdI (0.66 ± 0.16). VPS4a overexpression generated intermediate vesicle sizes (Z-average: 249 nm) with PdI of 0.80 ± 0.08. (J) Quantification of EV mean particle size (Z-average, nm). Tunicamycin treatment significantly altered vesicle size compared to control, while ESCRT modulation differentially affected vesicle diameter. (K) Polydispersity index (PdI) analysis showing changes in EV population heterogeneity across conditions. Tunicamycin increased vesicle heterogeneity, while ESCRT modulation produced variable effects on EV uniformity. Data in are presented as mean ± SEM with individual data points representing biological replicates (n = 6). Statistical analysis was performed using one-way ANOVA followed by Tukey’s post-hoc test for multiple comparisons. Significance is indicated as *p < 0.05, **p < 0.01, ****p < 0.0001; ns, not significant.

Modulation of ESCRT components revealed distinct functions for TSG101 and VPS4a. Knockdown of TSG101 (ShRNA TSG101) resulted in a concurrent reduction of both the exosomal marker CD63 and pathological TDP-43 (pTDP-43) (Fig. 6). This indicates that TSG101, a key component of ESCRT-I, is required for the general biogenesis and secretion of CD63-positive exosomes. In contrast, expression of a dominant-negative form of VPS4a (DN-VPS4a) specifically reduced levels of EV-associated pTDP-43 without affecting CD63 levels (Fig. 6). This demonstrates that the ATPase activity of VPS4a is dispensable for exosome secretion but is specifically required for the efficient loading of pTDP-43 cargo. Overexpression of wild-type TSG101 or wild-type VPS4a did not impair either CD63 or pTDP-43 levels in EVs compared to controls. These findings reveal a functional divergence within the ESCRT pathway: TSG101 is essential for core exosome formation, while VPS4a plays a specialised role in sorting pathological cargo into these vesicles.

#### Dynamic Light Scattering Identifies Distinct Vesicle Signatures Across ESCRT Conditions in a Tunicamycin-Induced MND Model

To characterise the physical properties of EVs isolated from primary neuron culture media under ESCRT modulation, we performed dynamic light scattering (DLS) analysis. The Z-average diameter (intensity-weighted mean hydrodynamic diameter) and polydispersity index (PdI) were measured for six independent EV preparations per condition (Figure 6). Under negative control (untreated) conditions, EVs exhibited a mean Z-average diameter of 247.7 ± 36.5 nm with a PdI of 0.44 ± 0.09, consistent with typical exosome-sized vesicles and moderate size distribution uniformity (Figure 6). Tunicamycin treatment (positive control) significantly reduced EV size to 112.5 ± 8.0 nm (p < 0.0001 vs. negative control) and substantially increased polydispersity (PdI = 0.94 ± 0.04), indicating that ER stress alters EV biogenesis and produces smaller, more heterogeneous vesicles (Figure 6F, G). Modulation of ESCRT components produced distinct effects on EV size and uniformity. TSG101 overexpression (+TSG101) resulted in significantly larger vesicles (386.2 ± 178.6 nm; p < 0.01 vs. negative control) with moderate polydispersity (PdI = 0.60 ± 0.23), suggesting enhanced exosome formation or altered cargo packaging (Figure 6G). In contrast, TSG101 knockdown (shTSG101) produced smaller, highly heterogeneous vesicles (138.7 ± 75.4 nm; PdI = 0.66 ± 0.16), consistent with disrupted exosome biogenesis (Figure 6I). VPS4a overexpression (+VPS4a) yielded EVs with Z-average diameter (248.9 ± 83.6 nm) and PdI (0.80 ± 0.08) comparable to negative controls, although with increased polydispersity (Figure 6F, G).

Representative size distribution profiles revealed that tunicamycin-treated and TSG101-knockdown samples exhibited broader, multi-modal distributions compared to the relatively monodisperse populations observed in negative control and TSG101-overexpressing samples (Figure 6E). Collectively, these DLS data demonstrate that tunicamycin-induced ER stress and ESCRT modulation significantly alter EV size and uniformity, complementing our Western blot findings (Figure 6) and further supporting the functional roles of TSG101 and VPS4a in exosome biogenesis and cargo sorting.

## Discussion

This study provides the first systematic characterisation of ESCRT protein expression in postmortem MND motor cortex and spinal cord, and demonstrates that modulating key ESCRT components such as TSG101, CHMP2B, and VPS4a profoundly influences TDP-43 pathology, autophagy, membrane protein stability, and exosome biogenesis. Collectively, these findings establish ESCRT dysfunction as a multifaceted contributor to MND pathogenesis and highlight ESCRT component’s critical role in neuronal homeostasis.

### ESCRT proteins are dysregulated in MND human tissues

Significant alterations in the expression of ESCRT proteins in sporadic MND patient brains and spinal cords are reported here for the first time. Specifically, CHMP2B (ESCRT-III) was significantly increased, while TSG101 (ESCRT-I) was significantly decreased in MND motor cortex compared to age-matched controls. These findings are consistent with previous reports linking CHMP2B mutations to frontotemporal dementia and MND [22, 23], and TSG101 dysfunction to impaired protein clearance in Alzheimer’s disease models [24, 25]. The reciprocal expression changes observed, upregulation of CHMP2B and downregulation of TSG101 suggest a potential compensatory or maladaptive response to proteostatic stress in MND. Notably, VPS37A, an ESCRT-0 component necessary for autophagosome completion [19], was also significantly reduced in MND motor cortex, extending previous reports linking VPS37A mutations to acute idiopathic myelitis and other neurological disorders [20]. In contrast, ALIX and STAM1 showed no significant changes, suggesting that ESCRT dysregulation in MND is complex and subunit-specific.

### TSG101 and CHMP2B differentially regulate TDP-43 pathology

TDP-43 aggregation is a hallmark of MND [7, 8, 27], and elevated pTDP-43 levels in MND brains were confirmed in the postmortem human tissue. Using cellular models of tunicamycin-induced ER stress, TSG101 overexpression was shown to significantly reduce both total and phosphorylated TDP-43, while TSG101 knockdown exacerbated TDP-43 accumulation. These findings align with the established role of TSG101 in endosomal sorting and autophagic clearance of ubiquitinated proteins [20] and are consistent with reports that TSG101 depletion reduces APP delivery to lysosomes, elevating intracellular amyloid-β [24] and impairing Tau clearance [25]. In contrast, CHMP2B exhibited a more selective role. CHMP2B overexpression reduced total TDP-43 levels, CHMP2B knockdown selectively decreased pTDP-43 without altering total TDP-43. This is consistent with recent reports that CHMP2B regulates TDP-43 phosphorylation via casein kinase 1 (CK1) independently of canonical autophagy pathways [28]. Mechanistically, CHMP2B overexpression stabilises CK1 by reducing its ubiquitination and lysosomal degradation, thereby sustaining CK1 kinase activity and promoting TDP-43 hyperphosphorylation [28]. Our data support this model: CHMP2B knockdown reduced pTDP-43 without altering tTDP-43. Thus, TSG101 and CHMP2B exert distinct regulatory effects on TDP-43: TSG101 primarily controls TDP-43 abundance, while CHMP2B influences its pathological phosphorylation.

### ESCRT modulation alters mTOR signalling and autophagy flux

Autophagy is a principal route for clearance of aggregated proteins in MND [13, 29], and our findings directly implicate the ESCRT machinery in its regulation. TSG101 overexpression significantly reduced both total and phosphorylated mTOR, while TSG101 knockdown increased p-mTOR levels. This is consistent with previous studies showing that TSG101 facilitates rapamycin-induced autophagic flux [30] TSG101 knockout impairs autophagic flux [31]. The reciprocal relationship between TSG101 and mTOR activity suggests that TSG101 promotes TDP-43 clearance through mTOR inhibition and downstream autophagic induction, a mechanism of direct therapeutic relevance given the promising preclinical and early clinical data for rapamycin-based strategies in neurodegeneration [37–40]. Rapamycin has shown efficacy in fly and mouse models of Huntington’s disease [37] and is currently under evaluation in clinical trials for MND and Alzheimer’s disease [38–40], supporting the translational potential of this axis. Consistent with impaired autophagic flux, TSG101 overexpression reduced p62 accumulation while TSG101 knockdown elevated both p62 and LC3B-II. Elevated p62 has been associated with TDP-43 mislocalisation and aggregation in MND and FTD [32, 33], and TSG101 deficiency recapitulates this phenotype in our models. Notably, CHMP2B modulation showed similar trends, although its effects on pTDP-43 appeared independent of bulk autophagy, consistent with the CK1-mediated pathway described above [28]. Together, these data indicate that ESCRT proteins regulate TDP-43 homeostasis through complementary mTOR-dependent and mTOR-independent mechanisms, the latter operating via CK1-mediated phosphorylation independently of bulk autophagy. This mechanistic bifurcation has important implications: therapeutic strategies targeting TSG101 and CHMP2B need not be mutually exclusive and may simultaneously address distinct aspects of TDP-43 pathology and mTOR-independent mechanisms.

### TSG101 is required for CD9 stability and membrane protein homeostasis with implications for exosome biogenesis

Tetraspanins such as CD9 are critical regulators of membrane organisation, intercellular signalling, and exosome biogenesis, and their dysregulation has been implicated in neurodegeneration. CD9 is enriched on plasma membrane-derived ectosomes and small EVs, where it participates in vesicle formation, cargo selection, and target cell interactions [34, 35]. In the present study, TSG101 depletion dramatically reduced CD9 expression under both basal and ER stress conditions, while TSG101 overexpression rescued tunicamycin-induced CD9 loss. This demonstrates that TSG101 is both necessary and sufficient for maintaining CD9 protein levels. DN-VPS4a expression, which blocks ESCRT-III-mediated membrane scission, also reduced CD9 levels, indicating that proper ESCRT function is required for CD9 homeostasis. This is consistent with reports that VPS4 activity is required for the release of CD9-positive EVs, and that VPS4 inhibition increases the concentration of ESCRT-III proteins recovered in EVs, suggesting a role for ESCRT-III disassembly during EV fission [34].

These observations extend known ESCRT functions beyond protein degradation to encompass active maintenance of neuronal membrane protein stability, likely by diverting CD9 from lysosomal degradation towards intraluminal vesicle-mediated recycling [17, 18]. Given the emerging evidence that tetraspanin dysregulation contributes to neuronal vulnerability in MND, the regulation of membrane protein stability, potentially through sorting CD9 away from lysosomal degradation or promoting its recycling via intraluminal vesicles [17, 18]. Given that membrane protein dysfunction contributes to MND pathogenesis, this represents a previously underappreciated role for ESCRT proteins in maintaining neuronal membrane integrity.

### TSG101 and VPS4a play distinct roles in exosome biogenesis and pathological cargo loading

Exosomes are established mediators of intercellular communication in neurodegeneration, facilitating the propagation of misfolded proteins between neurons [21]. Western blot characterisation confirmed that EV preparations were enriched in canonical exosomal markers (ALIX, TSG101, VPS4a) and free of the ER contaminant calnexin, validating preparation integrity. Modulation of ESCRT components revealed a clear functional divergence: TSG101 knockdown reduced both CD63 and EV-associated pTDP-43, demonstrating a requirement for TSG101 in general exosome biogenesis, while DN-VPS4a selectively depleted EV-associated pTDP-43 without affecting CD63 levels—indicating that VPS4a ATPase activity is dispensable for exosome secretion per se but essential for loading pathological TDP-43 cargo (Figure 7). This is consistent with the established role of VPS4a in ESCRT-III disassembly and membrane scission [17] and raises the possibility that selective VPS4a inhibition could reduce neuron-to-neuron propagation of pathological TDP-43 without broadly impairing exosome-mediated communication (Figure 7).

**Figure 7.**
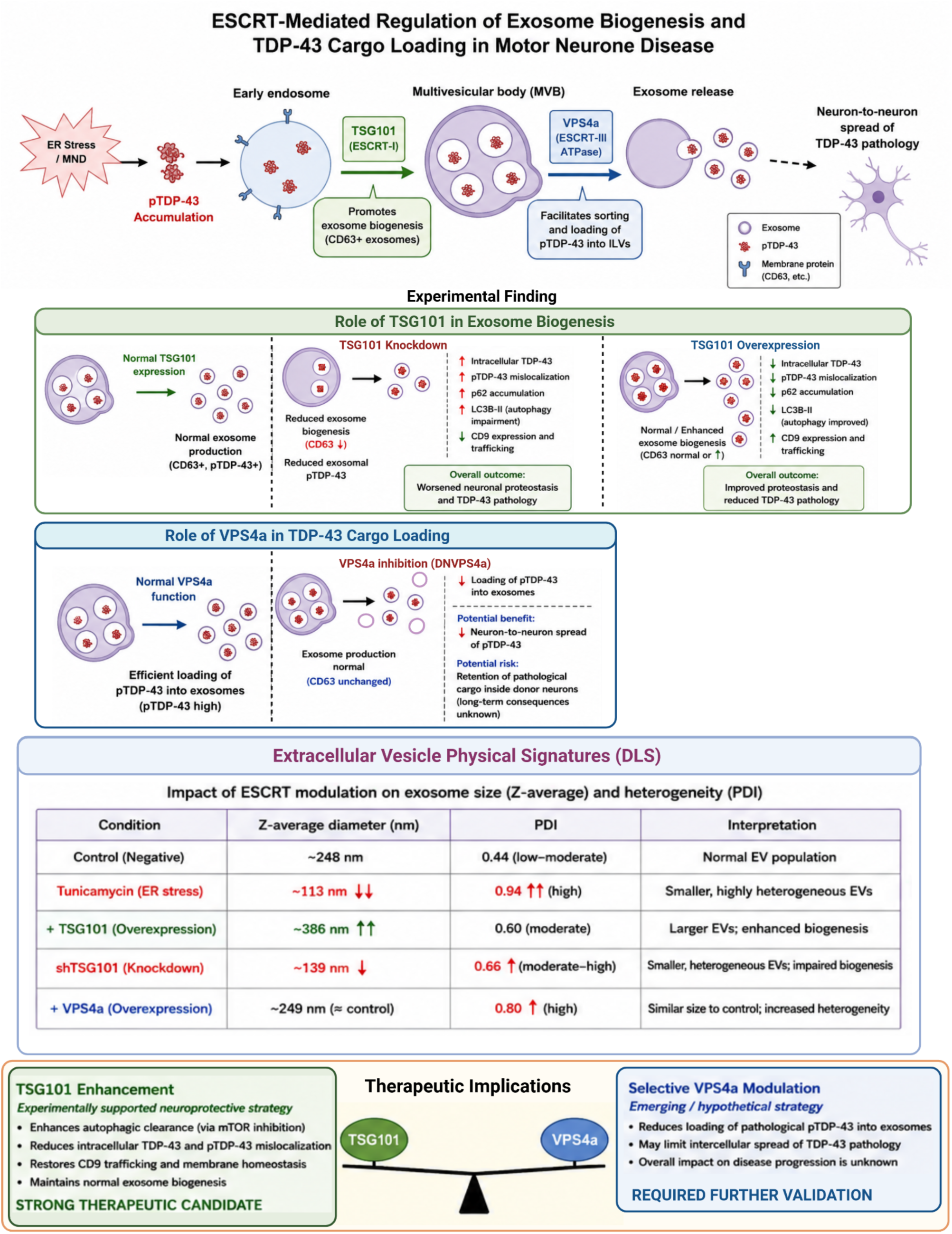
ESCRT-mediated regulation of exosome biogenesis and TDP-43 cargo loading in motor neurone disease (MND). Schematic overview of how ESCRT machinery components regulate extracellular vesicle (EV) biogenesis and pathological TDP-43 (pTDP-43) handling in a cellular model of MND. ER stress induces intracellular accumulation and mislocalisation of pTDP-43 at the early endosome, contributing to disease-associated proteostatic dysfunction. ESCRT-I component TSG101 promotes multivesicular body (MVB) formation and exosome biogenesis, particularly CD63-positive vesicles, thereby influencing extracellular export of pTDP-43. In contrast, ESCRT-III ATPase VPS4a regulates intraluminal vesicle (ILV) maturation and facilitates selective sorting and loading of pTDP-43 into exosomes, potentially modulating intercellular propagation of pathology. Experimental manipulations demonstrate differential roles of ESCRT components. TSG101 knockdown reduces exosome biogenesis (CD63-positive EVs), decreases extracellular pTDP-43 release, and is associated with impaired autophagic flux (p62 accumulation, LC3B-II dysregulation), altered CD9 trafficking, and increased intracellular pTDP-43 burden, collectively exacerbating neuronal proteostasis defects. Conversely, TSG101 overexpression enhances exosome production, improves autophagy markers, restores membrane trafficking homeostasis, and reduces intracellular pTDP-43 accumulation, supporting a neuroprotective role. Functional modulation of VPS4a indicates that inhibition (DNVPS4a) reduces pTDP-43 loading into exosomes while maintaining overall exosome release, suggesting a selective role in cargo sorting rather than vesicle biogenesis. However, this may limit extracellular spread of pathological TDP-43 with potential trade-offs in intracellular retention. Dynamic light scattering (DLS) analysis reveals that ESCRT modulation alters EV physical properties: ER stress (tunicamycin) produces smaller, highly heterogeneous vesicles (reduced Z-average, increased PDI), while TSG101 overexpression increases vesicle size and reduces heterogeneity. In contrast, TSG101 knockdown generates smaller, heterogeneous EVs, whereas VPS4a modulation minimally affects vesicle size but increases heterogeneity. Overall, the findings highlight a dual and context-dependent role of ESCRT machinery in regulating EV biogenesis, cargo sorting, and TDP-43 pathology propagation, with TSG101 emerging as a potential neuroprotective therapeutic target, while VPS4a represents a candidate for selective modulation of pathological cargo dissemination.

DLS characterisation provides additional biophysical evidence that ESCRT modulation influences extracellular vesicle (EV) biogenesis. ER stress induced by tunicamycin produced smaller and more heterogeneous EV populations, suggesting that proteostatic stress disrupts normal multivesicular body maturation and vesicle formation. In contrast, TSG101 overexpression increased EV size, consistent with enhanced intraluminal vesicle formation and more efficient exosome biogenesis, whereas TSG101 depletion generated smaller, more heterogeneous vesicles, indicative of impaired ESCRT-dependent membrane remodeling (Figure 7). These observations are consistent with previous studies demonstrating that disruption of ESCRT function alters EV morphology, size distribution, and cargo composition, reflecting defective multivesicular body dynamics and intraluminal vesicle formation [59, 60]. Together with our biochemical analyses, these findings support a central role for TSG101 in regulating both the physical characteristics of EVs and the efficient biogenesis of CD63-positive exosomes, whereas VPS4a appears to primarily regulate the selective incorporation of pathological TDP-43 cargo rather than overall vesicle production (Figure 7).

### Therapeutic implications and future directions

The present findings identify TSG101, CHMP2B, and VPS4a as mechanistically distinct therapeutic targets in MND. Restoring TSG101 expression or function could promote TDP-43 clearance via enhanced autophagy and maintain membrane protein homeostasis. Conversely, modulating CHMP2B activity might reduce pathological TDP-43 phosphorylation without affecting total TDP-43 levels, offering a strategy to mitigate TDP-43 toxicity while preserving its nuclear functions. However, several limitations must be acknowledged. The human tissue sample size (n = 6 per group) is relatively small, and findings should be validated in larger independent cohorts. In addition, the mechanistic studies were conducted in cell lines and primary neurons; *in vivo* validation in animal models of MND (e.g., TDP-43 transgenic) is needed to establish causal relationships. The precise molecular mechanisms by which TSG101 and CHMP2B regulate TDP-43 phosphorylation and autophagy require further investigation, including identification of direct interaction partners.

Nevertheless, the convergence of evidence from human tissues, cellular models, and biophysical characterisation strongly supports a central role for ESCRT dysfunction in MND pathogenesis. Future studies should explore whether pharmacological enhancement of TSG101 activity or selective inhibition of CHMP2B-mediated TDP-43 phosphorylation can slow disease progression in preclinical models. Additionally, the distinct roles of TSG101 and VPS4a in exosome cargo loading suggest that targeting these proteins could modulate the spread of pathological TDP-43 between neurons, an increasingly recognised mechanism in MND progression.

## Conclusion

In summary, this study is the first to characterise ESCRT protein expression in human MND tissues and to define the mechanistic consequences of ESCRT modulation across multiple disease-relevant pathways. TSG101 and CHMP2B exert distinct, complementary effects on TDP-43 pathology: TSG101 promotes clearance through mTOR-dependent autophagic induction, while CHMP2B selectively modulates pathological phosphorylation via a CK1-dependent, autophagy-independent mechanism. In parallel, TSG101 and VPS4a perform divergent functions in exosome biogenesis—TSG101 supporting general vesicle production, VPS4a mediating selective loading of pathological TDP-43 cargo. These mechanistically distinct roles open complementary therapeutic avenues: restoring TSG101 function to enhance proteostatic clearance, inhibiting CHMP2B–CK1 signalling to reduce TDP-43 hyperphosphorylation and targeting VPS4a to limit prion-like intercellular spread of pathological TDP-43. Validating these strategies *in vivo* represents a critical next step towards disease-modifying treatments for MND.

## Funding and Acknowledgements

This work was supported by the Ministry of Higher Education of the Arab Republic of Egypt (PhD scholarship to L.M.). We thank London Neurodegenerative Diseases Brain Bank for sharing the post-mortem brain tissue critical for this study.

## Author Contributions

L.M. and M.F.S. contributed equally to this work. Conceptualisation: S.K.; Methodology: L.M., M.F.S., R.W., S.L.M., S.K.; Formal analysis and investigation: L.M., M.F.S. SK; Writing original draft preparation: L.M., M.F.S.; Writing, review and editing: R.W., S.L.M., S.K.; Supervision: S.K.; Funding acquisition: L.M., M.F.S., SK. All authors read and approved the final manuscript.

## Ethics Approval

Human postmortem tissue use was approved by the relevant institutional review boards of the London Neurodegenerative Diseases Brain Bank, Institute of Psychiatry, Psychology and Neuroscience, King’s College London (London, UK). The Brain Bank operates under Human Tissue Authority (HTA) licensing and receives generic ethical approval from the London - City & East Research Ethics Committee (REC reference 18/WA/0206, date of favourable opinion 27 June 2018). This approval covers the release of anonymised tissue to external researchers for approved projects, obviating the need for project-specific REC review. All donors or their nominated next-of-kin provided written informed consent for the collection, storage, and use of tissue in research. Animal procedures were approved by the University of Bradford Animal Welfare and Ethical Review Body and performed in accordance with the UK Animals (Scientific Procedures) Act 1986.

## Consent to Participate

All tissue donors or their next of kin provided written informed consent for the use of biological material in research.

## Consent to Publish

Not applicable. This study does not contain data from identifiable individual participants.

## Data Availability

The datasets generated and/or analysed during the current study are available from the corresponding author (S.K.) on reasonable request.

